# Megafauna decline have reduced pathogen dispersal which may have increased emergent infectious diseases

**DOI:** 10.1101/2020.01.21.914531

**Authors:** Christopher E. Doughty, Tomos Prys-Jones, Søren Faurby, Andrew Abraham, Crystal Hepp, Victor Leshyk, Viacheslav Y. Fofanov, Nathan C. Nieto, Jens-Christian Svenning, Mauro Galetti

## Abstract

The Late Quaternary extinctions of megafauna (defined as animal species >44.5 kg) reduced the dispersal of seeds and nutrients, and likely also microbes and parasites. Here we use body-mass based scaling and range maps for extinct and extant mammal species to show that these extinctions led to an almost seven-fold reduction in the movement of gut-transported microbes, such as *Escherichia coli* (3.3 km^2^/day to 0.5 km^2^/day). Similarly, the extinctions led to a seven-fold reduction in the mean home ranges of vector-borne pathogens (7.8 km^2^ to 1.1 km^2^). To understand the impact of this, we created an individual-based model where an order of magnitude decrease in home range increased maximum aggregated microbial mutations 4-fold after 20,000 years. We hypothesize that pathogen speciation and hence endemism increased with isolation, as global dispersal distances decreased through a mechanism similar to the theory of island biogeography. To investigate if such an effect could be found, we analysed where 145 zoonotic diseases have emerged in human populations and found quantitative estimates of reduced dispersal of ectoparasites and fecal pathogens significantly improved our ability to predict the locations of outbreaks (increasing variance explained by 8%). There are limitations to this analysis which we discuss in detail, but if further studies support these results, they broadly suggest that reduced pathogen dispersal following megafauna extinctions may have increased the emergence of zoonotic pathogens moving into human populations.

## Introduction

Since the Late Pleistocene and early Holocene, the loss of the planet’s largest mammals has affected trophic structure, seed dispersal, biogeochemistry and nutrient dispersal globally (Galetti et al., 2018) (Malhi et al., 2016) (Doughty et al., 2016). Large animals play unique roles in dispersal processes because their long gut lengths, daily movements, and home ranges enable them to carry seeds, spores and nutrients long distances across landscapes. Obligate ectoparasites (such as ticks, fleas, lice) and microbes residing in the gut or other tissues rely on animal hosts for transport. Following the megafauna extinctions, the mean dispersal distance of both would have been reduced. Could such changes in microbe dispersal distances have had broader ecosystem consequences? Here, we ask whether changes in microbe dispersal distances as a consequence of the megafauna extinctions impacted the emergence of zoonotic infectious diseases.

Understanding the predictors of zoonotic emergent infectious disease (EIDs) will enable better prediction, surveillance and management of future disease outbreaks. A study by Jones et al (2008) aggregated 335 EID origin events between 1940 and 2004, and found that 60.3% are zoonoses, with over 71.8% of these having a wildlife origin (n = 145) (Jones et al., 2008). The authors found that host species richness was a significant predictor of zoonotic pathogens emerging from wildlife populations. Other studies have shown that infectious diseases emerged through humanity’s close association with agriculture and domestic animals (Dobson & Carper, 1996) (Wolfe, Dunavan, & Diamond, 2007). A closer proximity with animals and higher human population densities increased the establishment and spread of EIDs. Infectious diseases existed in hunter-gatherers but were subject to differing evolutionary pressures that allowed them to persist in low population densities (versus high population densities of agrarian societies). A fast-acting, highly virulent disease would quickly kill off the sparse hunter-gather population before the disease had a chance to spread, thus also killing off the disease.

The rise of agriculture and settling of peoples into close-knit communities clearly impacted disease emergence, but could the Pleistocene and early Holocene megafauna extinctions (Sandom et al 2014) also have shaped infectious disease? There is strong debate about whether there is a positive biodiversity disease relationship especially related to human pathogens. Biodiversity loss tends to increase disease occurrence because the lost species are initially replaced with more abundant generalists that invest more in growth and less in adaptive immunity(Keesing et al., 2010), making them better hosts for pathogens. Additionally, biodiversity loss may increase disease occurrence due to a reduction in the dilution effect. This posits that biodiversity decreases the probability of an outbreak by diluting the assemblage of transmission-competent hosts with non-competent hosts (Schmidt and Ostfeld 2001). However, others have found that increases in biodiversity over time were not correlated with improved human health (Wood et al., 2017). Here, our research focuses on how EIDs might have been affected by the loss of dispersal, which has been decreasing through large animal population declines, extinctions, and more recently through human restrictions including fences and roads (Doughty et al., 2016) (Tucker et al., 2018). We hypothesize that as global dispersal distances decreased following the megafauna extinctions, pathogen speciation may have increased with isolation in a mechanism similar to the theory of island biogeography(Reperant, 2010) (Heaney, 2000). This reduced movement may also impact EID formation by increasing the immune-naivety of the remaining host species because they will no longer regularly interact with as many pathogens.

In this paper, we first quantify the global change in pathogen dispersal through faeces (dispersed through the gut) and obligate ectoparasites (e.g. ticks) before and after the Late Pleistocene/early Holocene megafauna extinctions. Next, we create an individual-based model to mechanistically show how reduced dispersal could impact aggregated microbial genetic change over time. Finally, we test whether the global decrease in pathogen dispersal impacted EID formation. If the loss of dispersal is important to EID formation, then the regions of greatest dispersal loss will be statistically correlated to EID formation. Previous papers have statistically correlated EID outbreaks with human population density, mammal biodiversity, rainfall and found statistical patterns strong enough to make predictions about potential future outbreaks (Jones et al., 2008). We then add changes to dispersal patterns over time to see if the prediction of EIDs is improved.

## Materials and Methods

### I. Impact of megafauna extinctions on microbial and blood parasite movement

We estimated current global ecto-parasite and fecal pathogen dispersal patterns using the IUCN mammal species range maps for all extant species (removing all bats because mass scaling of dispersal for these taxa is inaccurate) (N=5,487). To create maps of dispersal patters for a world without the Pleistocene megafauna extinctions, we added species range maps (N=274) of the now extinct megafauna (within 130,000 years) created in Faurby & Svenning (2015a) to the current IUCN based dispersal maps. These ranges estimate the natural range as the area that a given species would occupy under the present climate, without anthropogenic interference. In cases of evident anthropogenic range reductions for extant mammals, like the Asian elephant (*Elephas maximus*), the current ranges encompass only the IUCN defined ranges. However, our models of the world without extinctions includes the ranges on these extant animals prior to anthropogenic range reductions. The taxonomy of recent species followed IUCN while the taxonomy of extinct species (which were included if there are dates records less than 130,000 years old) followed Faurby & Svenning (2015b). Each living and extinct animal species was assigned a body mass estimate (Faurby & Svenning, 2016), with the few species lacking these estimates being assigned masses based on the mass of their closest relatives. We used the following mass based (M: average body mass per species (kg)) scaling equations (recalculated from primary data in Figure S1) to estimate home range (Kelt and Van Vuren, 2001), day range (Carbone et al 2005), and gut retention time (Demment and Van Soest, 1985, Demment 1983):

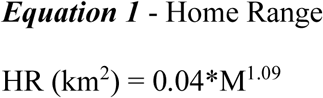

This dataset, originally compiled by Kelt and Van Vuren (2001) (N=113 mammalian herbivores), used the convex hull approach to calculate home range and found the mass-based scaling to be highly size dependent (with mass scaling exponent of >1)

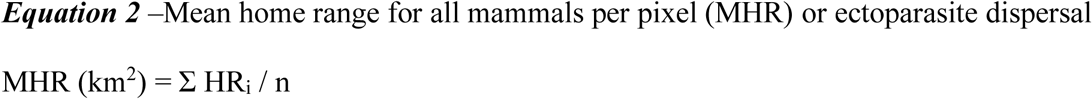

(i: per pixel species number; n: = number of mammal species per pixel)

We define the mean ectoparasite dispersal per pixel as the average distance a pathogen could travel across all mammals present in the pixel and assuming an equal change of colonizing any mammal species.

Next, we estimate fecal pathogen diffusivity with the following equations. We start with day range (daily distance travelled) originally from Carbone et al 2005 (N=171 mammalian herbivores) but recalculated from primary data in Figure S1.

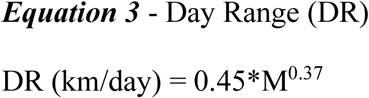

Next, to estimate the minimum time a generalist microbe might stay in the body of a mammalian herbivore, we use passage time:

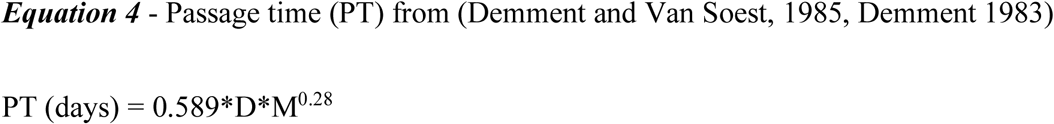

Where D is digestibility, which we set to 0.5 as a parsimonious assumption because the actual value is unknown for many extant and extinct animals.

Distance between consumption and defecation or straight line fecal transmission distance is simply multiplying equation 3 by equation 4:

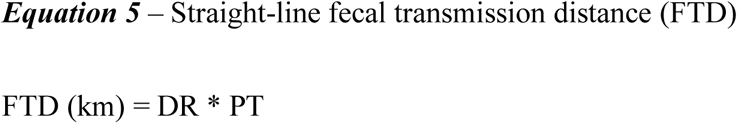

However, animals rarely move in a straight line, and without any additional information, we can assume a random walk pattern with a probability density function governed by a random walk as:

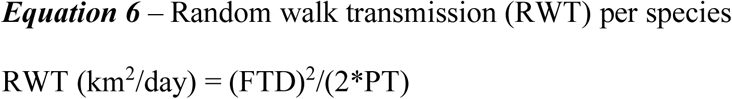

Here we define the mean fecal diffusivity as the mean range in any pixel a generalist microbe could travel during its lifetime assuming an equal chance of colonizing any mammal species.

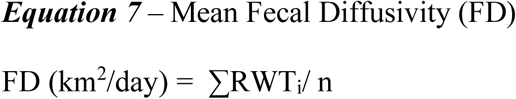

(i: per pixel species number; n: = number of mammal species per pixel)

This equation represents the average distance a fecal pathogen would travel in an ecosystem if it had an equal chance of being picked up by any nearby species walking in a random walk.

### Individual based model

To establish whether the loss of terrestrial megafauna increased microbe heterogeneity, we used Matlab (Mathworks) to create an individual based model (IBM) with two randomly distributed animal species carrying a generalist microbe. We varied our model assumptions (parentheses below) in sensitivity studies (Tables S7). The IBM consisted of a 500×500 cell grid (300×300 and 1000×1000 – in our sensitivity study, we tried big and small grids) with species A in 10% (5 and 20%) of randomly selected cells and species B also in 10% (5 and 20%) of cells. 10% (5 and 20%) of animals contained the generalist microbe. We then created a 9 by 9 grid around each of species A. This was considered the home range of the species and the group of animals would interact with all other groups of animals within that home range. We assumed the home range of species B to be a single grid cell. We make a simple assumption that mutations in this generalist microbe increase linearly with time until two animals interact, at which point the microbe is assumed to have been shared and the accumulated difference between host microbiomes is reset to zero. Later, we reduced the home range of species A from a 9×9 to a 3×3 grid, mimicking the decline in dispersal following the extinctions. We then, at each time step, identified the microbe with the highest number of accumulated mutations within the 500 by 500 grid for the megafauna world (9 by 9 simulation) and the post extinction world (3 by 3 simulations) (Figure 2). In order to parameterize the model with real world values, we assigned a single time step an arbitrary value of a single year (see justification in supplementary methods). The model was run for 20,000 years, putting the range reduction of species A at around 10,000 years ago, an approximate date for a large part of the Late Pleistocene extinctions.

**Figure 1.**
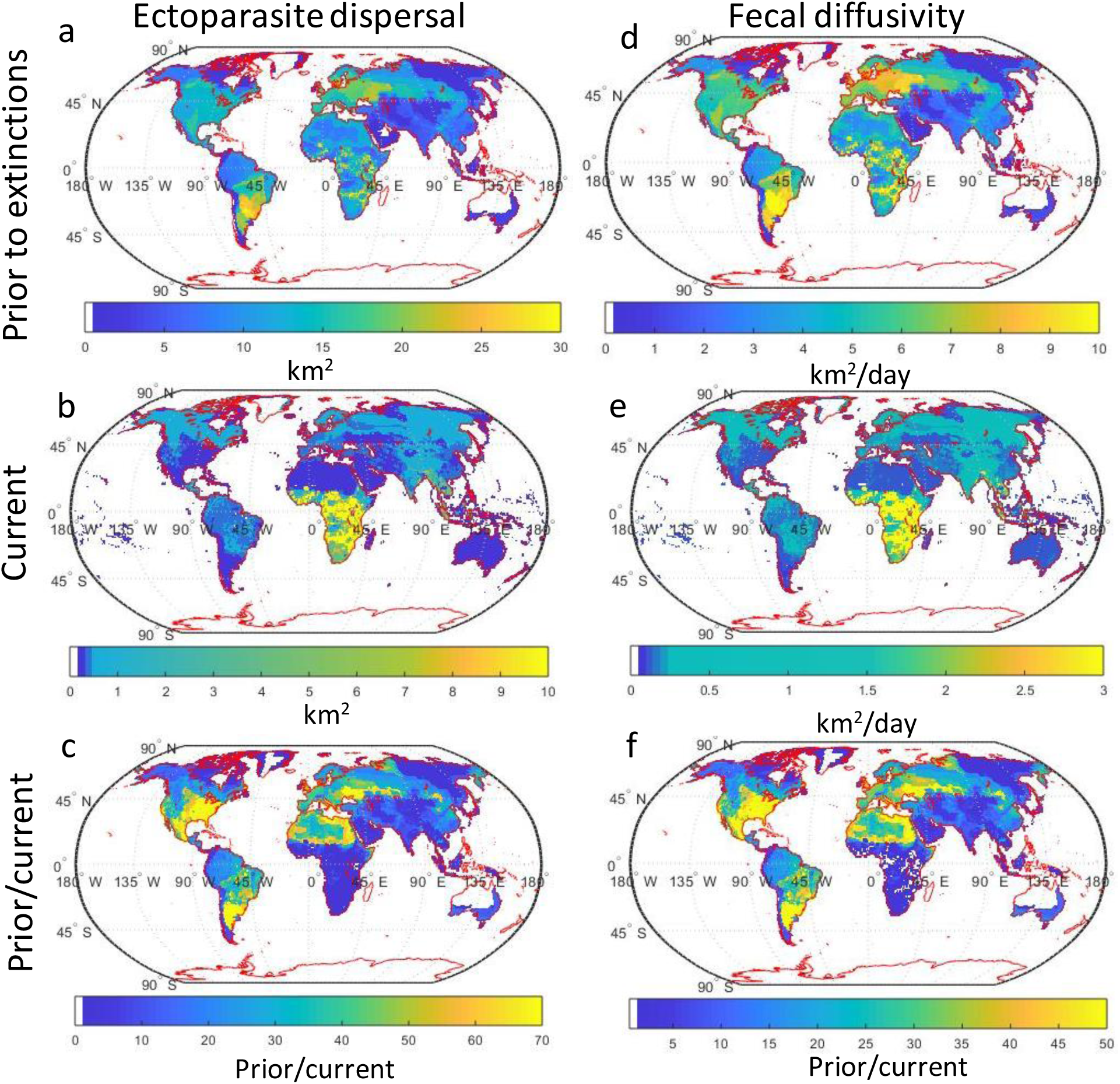
Maps of (left) mean ectoparasite dispersal (km^2^) (from equation 2) and (right) fecal diffusivity (km^2^/day) (from equation 7) for the world prior the megafauna extinctions (top), with current animals (middle), and prior divided by current (bottom).

**Figure 2.**
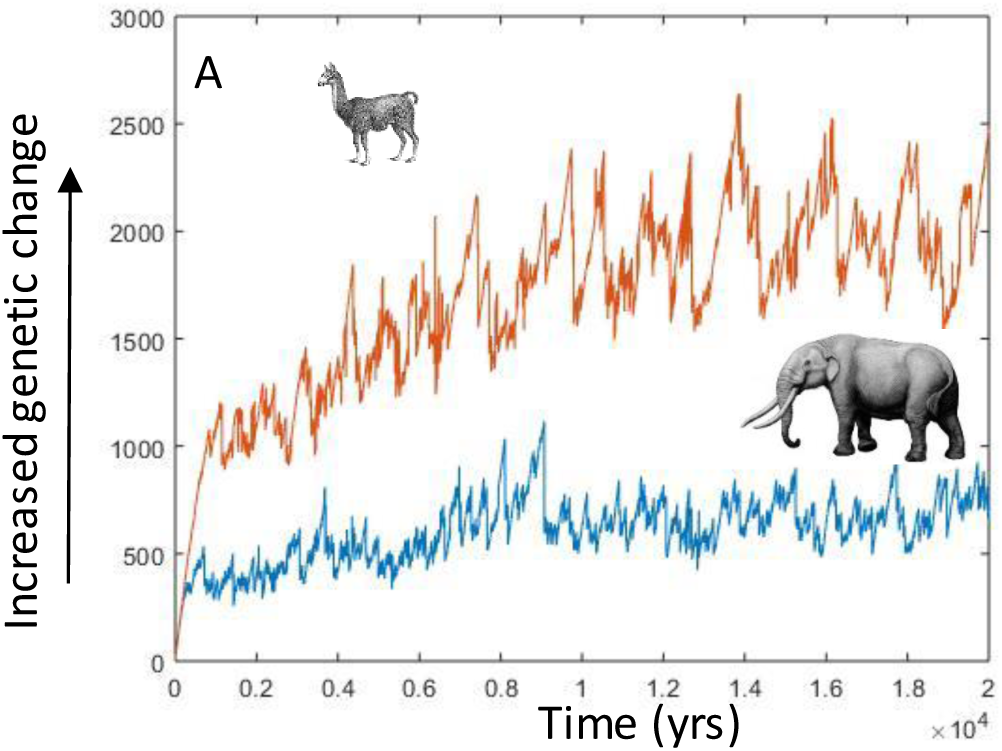
The maximum aggregated genetic changes per grid (500 by 500) per time step when animals were constrained to move within a 9×9 pixel space (blue – megafauna world represented by the Stegomastodon), and a 3 by 3 pixel space (red – current world represented by the llama). A sensitivity study for our parameters is show in Figure S7.

### EID modelling

We then tested whether these changes in pathogen dispersal distance could help explain the location of 145 new zoonotic diseases (with a wildlife origin) that emerged over the past 64 years (Jones et al., 2008). Jones et al 2008 searched the literature to find biological, temporal and spatial data on 335 human EID ‘events’ between 1940 and 2004 of which 145 were defined as zoonotic. We also divide our analysis into vector driven (Table S3), non-vector driven (Table S4) and all diseases (Table 2). To control for spatial reporting bias, they estimated the mean annual per country publication rate of the Journal of Infectious Disease (JID). However, this is not a perfect control for reporting bias as it may bias towards first world countries. In their paper, they used predictor variables of log(JID), log(human population density), human population growth rates, mean monthly rainfall, mammal biodiversity, and latitude. We repeat this study but add six data layers shown in Figure 1 of animal function, as well as other variables such as rainfall seasonality, total biodiversity (species richness including the now extinct megafauna), biomass weighted species richness and the change in biomass weighted species richness. In total, we tested 16 variables against the EID outbreaks (explained in Table S1). In addition to the 145 known EID outbreaks, we randomly generated ∼five times more random points (>600 points) to compare them (all results in the paper are the average of three separate runs where the control points vary randomly) (see Figure S2 as an example distribution).

**Table 1.**
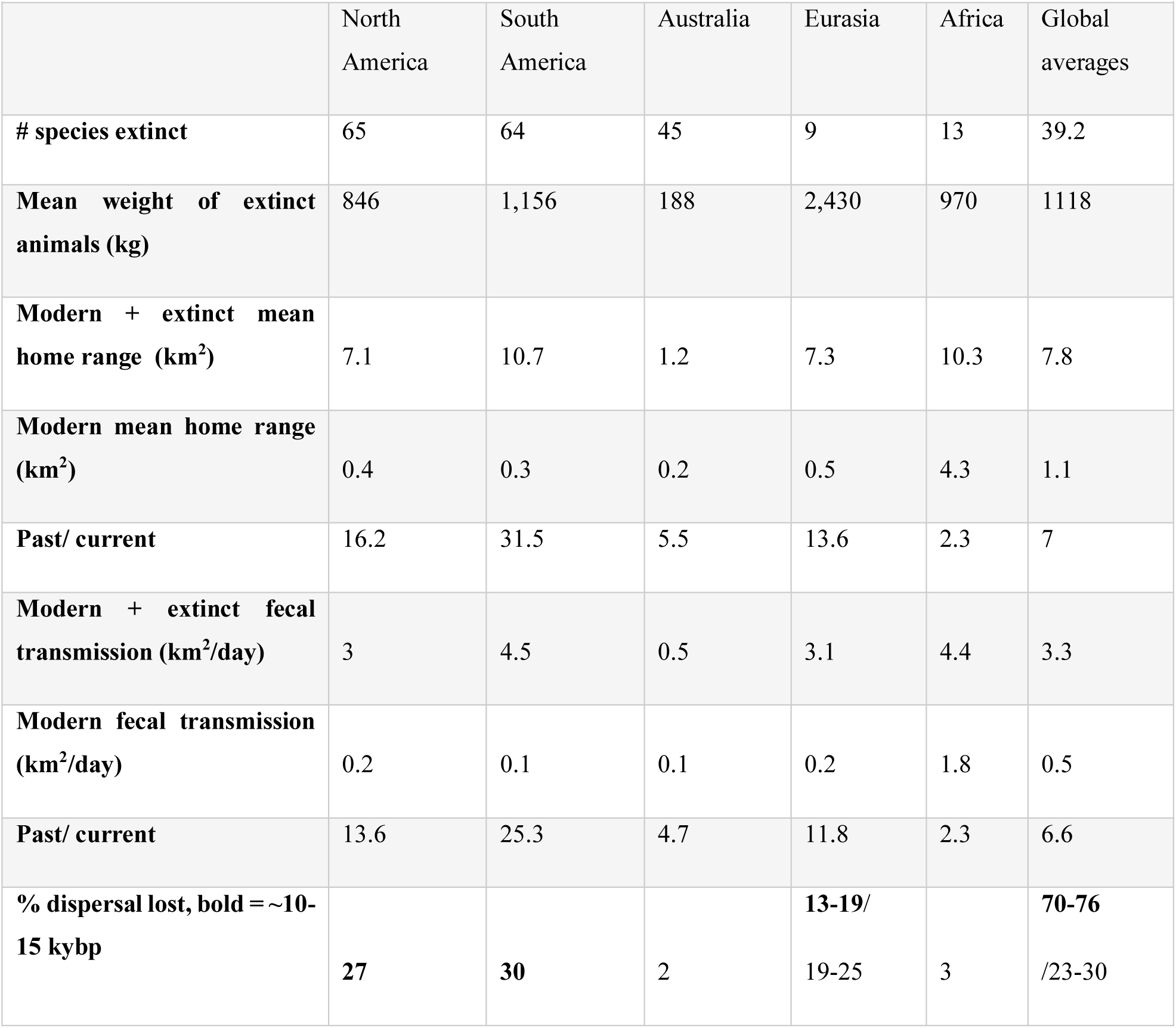
The number of animal species that went extinct between the Late Pleistocene and early Holocene, their mean weight, average home range (km^2^) and fecal transmission (km^2^/day) for each continent and the globe calculated for modern species and for modern plus extinct species. Also shown is the change (past/current) between these periods calculated for each pixel at the global scale. % dispersal lost is the lost continental dispersal divided by total lost global dispersal (weighted by area and excluding Antarctica). Bold numbers represent dispersal lost ∼10-15 kybp and the range for Eurasia represents North to South differences and uncertainty (33-50%) in Eurasian extinctions and not bold represents other parts of the world.

**Table 2.** A SAR^err^ analysis to predict the presence of 145 EIDs compared with ∼600 randomly generated points using 16 variables described in Table S1. Using AIC, r^2^ and VIF we show the four best models with the predictors of *JID* -journal of infectious disease articles, human population density, *SR*_*current*_ current species richness, *Δ MHR* – change in mean home range (Figure 1c and eq 2), *ΔFD* – change in fecal diffusivity (Figure 1f and eq 7), and Rain – average rainfall. In column one, we show the variables of interest, in column two, we show the individual model coefficient, r^2^ and significance using the Bonferroni correction to determine significance or *0*.*05/16= 0*.*003125 =* * (*=P<*0*.*003125*, \*\**=* P<*06e-4*, \*\*\**=* P<*06e-5*). For the rest of the models, we use standard significance (*=P<*0*.*05)*.

| Variable | Individual | Model null -<br>pseudo $r^2 = 0.185$,<br>AIC=460 | Model FD -<br>pseudo $r^2 =$<br>**0.201, AIC=448** | Model HR -<br>pseudo $r^2 =$<br>**0.197, AIC=452** | Model SR -<br>pseudo $r^2 =$<br>**0.201, AIC=450** |
| --- | --- | --- | --- | --- | --- |
| **(Intercept)** | | $-0.08 \pm 0.026$ *** | $-0.12 \pm 0.027$ *** | $-0.11 \pm 0.027$ *** | $-0.11 \pm 0.05$ |
| **Log(*JID*)** | $r^2 = 0.046$ ,<br>0.065 *** | $0.072 \pm 0.01$ *** | $0.064 \pm 0.01$ *** | $0.065 \pm 0.01$ *** | $0.063 \pm 0.01$ *** |
| **Pop density** | $r^2 = 0.12$ , 0.14<br>*** | $0.13 \pm 0.014$ *** | $0.13 \pm 0.014$ *** | $0.13 \pm 0.014$ *** | $0.13 \pm 0.014$ *** |
| **$\Delta FD$ (eq7)** | $r^2 = 0.042$ ,<br>0.004 *** | | $0.0026 \pm 0.0007$<br>*** | | $0.0026 \pm 0.0007$<br>*** |
| **$\Delta MHR$ (eq2)** | $r^2 = 0.035$ ,<br>0.003 *** | | | $0.0017 \pm 0.0005$<br>*** | |
| **$SR_{current}$** | $r^2 = 0.012$ ,<br>0.001 * | | | | $8e-6 \pm 5e-4$ |
| **Rain** | $r^2 = 0.021$ ,<br>0.0009 ** | $0.0004 \pm 0.0002$ * | $0.0004 \pm 0.0002$ * | $0.0004 \pm 0.0002$ * | $0.0004 \pm 0.003$ |

We then used the Ordinary Least Squares (OLS) multiple regression models to predict the EID events. We used Akaike’s Information Criterion (AIC) for model inter-comparison, corrected for small sample size. Whenever spatial data are used there is a risk of autocorrelation because points closer to each other will have more similar signals than points far from each other. We therefore used Simultaneous Auto-Regressive (SAR^err^) models (Table 2) to account for spatial autocorrelation (Dormann et al., 2007) using the R library ‘spdep’ (Bivand, Hauke, & Kossowski, 2013). SAR-err reduces the sample size by assuming that all outbreaks within the same neighbourhood are the same. We examined possible neighbourhood sizes to determine how effective each was at removing residual autocorrelation from model predictions. We defined neighbourhoods by distance to the sample site. We tried distances from 5 km to 300 km and found that AIC was minimized at 200 km (Figure S3 – the average of 16 simulations). Following this reduction of our dataset, our correlogram (Figure S4) indicates vastly reduced spatial autocorrelation. We estimated the overall SAR model performance by calculating the square of the correlation between the predicted (only the predictor and not the spatial parts) and the raw values. We will refer to this as pseudo-R^2^ in the paper even though we are aware that several different estimates of model fit are frequently referred to as pseudo-R^2^. We also did a VIF analysis using the R package usdm (Naimi et al., 2014) to control for multicollinearity and all VIFs of the predictor variables are below 1.5 showing little multicollinearity.

## Results

Without the megafauna extinctions, we estimate that the mean global home range of a generalist ectoparasite (Equation 2 – the average home range of all (non-bat) mammal species in its ecosystem) averages ∼8 km^2^ (Table 1). In parts of Eurasia and southern South America, that had a particularly high pre-extinction diversity of large-mammals, the mean home range exceeded 25 km^2^ (Figure 1). Following the extinctions, the mean global home range of a generalist blood parasite has been reduced to 1.1 km^2^ or 14% of the previous global average (Table 1). The decreases were particularly large in South America, where mean home range decreased over two orders of magnitude in the south of the continent since all 26 local mammal species >1,000 kg went extinct (Sandom et al 2014).

The mean fecal diffusivity (Equation 7) is the minimum distance (assuming microbes are excreted in the first defecation) a generalist gut pathogen could travel between consumption and defecation and is highly size dependent. Without megafauna extinctions, this area is greater than 3.3 km^2^/day and up to 10 km^2^ /day in parts of Eurasia and southern South America (Figure 1). Outside of abiotic dispersal by wind or water, this is a potentially important way for microbes to move across an ecosystem. Following the megafauna extinctions, the mean distance travelled by microbes globally through biotic means decreased to 0.5 km^2^ or ∼15% of the non-extinction value. The largest declines in distance travelled are in the Americas and Eurasia.

We estimated the approximate increase in time for fecal pathogens and obligate ectoparasites to travel the same distance by dividing maps without extinctions by current maps (180×360) of mean ectoparasite and fecal dispersal (Figure 1). For instance, in southern South America, it would take 70 times as long for ectoparasites and 50 times as long for fecal microbes to travel the same distance without versus with megafauna (Figure 1c and f). This contrasts to parts of Africa, where there is little change.

To better understand the possible impacts of lost dispersal on microbes, we used an individual based model (IBM) where initially, a large animal with a large home range periodically shared microbes with a small animal with a small home range. We make a simple assumption that mutations in this generalist microbe increase linearly with time until two animals interact, at which point the microbe was assumed to have been shared and the mutational difference counters for each were reset to zero. Upon replacing an animal with a large home range (81 pixels) with a smaller home range (9 pixels), after 20,000 years of simulation time (Figure 2 and SI Appendix, Table S7), the maximum aggregate mutations increased fourfold in the 3 by 3 (compared to the 9 by 9), but did not saturate for >10,000 years illustrating why it is important to understand EID drivers over long timescales. Although vastly oversimplified, our model demonstrates that long periods of time (∼10,000 years) might be necessary for genetic changes to build up following the loss of dispersal. Most of the global dispersal capacity (70-76% - Table 1) was lost at this timescale (∼10,000 years; and most of the loss before this time point was concentrated in Australia) near the end of the last ice age ∼10-15 kybp following extinctions of megafauna from North and South America and Siberia.

How might we theoretically predict reductions in microbe and ectoparasite dispersal to impact pathogen formation? We would first predict that an extinct animal like a mammoth or mastodon would have a large home range (red line Figure 3a) because home range is highly size-dependent (Wolf et al., 2013) and isotope data suggest this is true (Hoppe et al., 1999). Such a large animal would also host many ectoparasite species because parasite species richness is also size dependent (Esser et al., 2016). Paleodata shows the extinct megafauna did indeed host parasites (McConnell & Zavanda, 2013). In our qualitative example for South America (Figure 3), the *Stegomastodon* home range overlaps with species of extant mammals, vectors and likely microbes/pathogens. Home range overlap between the species would ensure periodic interactions with pathogens so all would develop some immune protection. In Figure 3b, the Stegomastodon goes extinct and the smaller species no longer regularly interacts with other pathogens, becoming immune-naïve and more susceptible to generalist pathogens when they eventually interact. This isolation would allow variation to accumulate in the microbes, leading to possible evolutionary divergence (represented by several colors of the pathogens in Figure 3b). Figure 3c shows late Holocene interactions with humans and their immuno-limited domestic ungulates (inbred animals are more susceptible to pathogens) (Smallbone, van Oosterhout, & Cable, 2016), highlighting other important variables necessary for EID occurrence(Jones et al., 2008, Table 2). The arrival of domestic animals is thought to be important for EID occurrence because there is an evolutionary trough that needs to be surpassed for a pathogen to colonize a new host and evidence suggests this may be lower in domestic animals (Smallbone et al., 2016).

**Figure 3.**
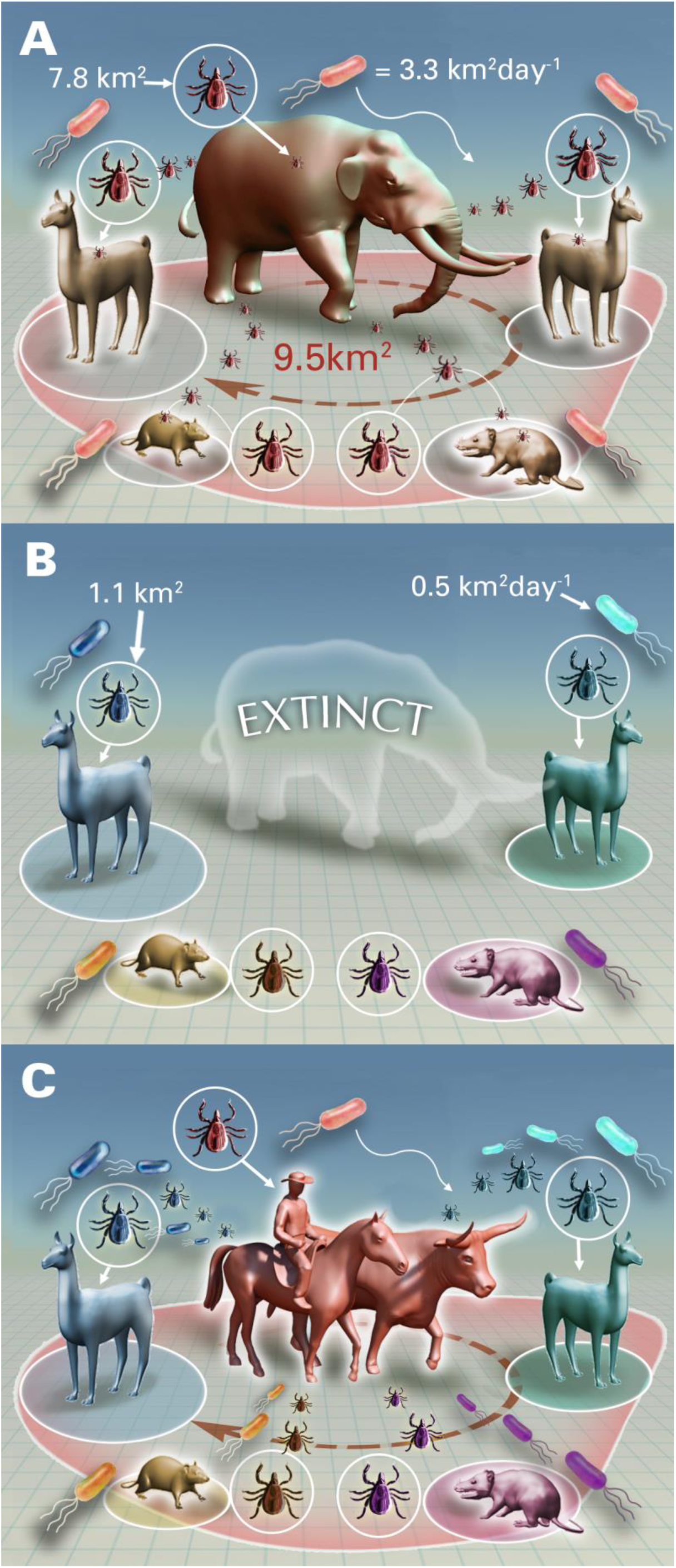
(A) Hypothetical example of a South American late Pleistocene animal assemblage and their home ranges. The animals host tick-borne and fecal pathogens with homogenous colors because the large home range of the megafauna keeps them interacting. Numbers indicates the mean global dispersal distance (Table 1) for ectoparasites and fecal pathogens. (B) Hypothetical early to middle Holocene animal assemblage without the extinct megafauna, thus removing the regular interaction between tick and fecal pathogens increasing immuno-naivety for all species. Colors of tick-borne and fecal pathogens begin to diverge representing a hypothetical speciation because without the now extinct megafauna there is less interaction between pathogens. (C) Hypothetical late Holocene animal assemblage with humans and their domestic animals picking up the now diverged (many colored) pathogens which could cause EIDs in people and domestic animals. This panel has no numbers because we did not calculate Anthropocene pathogen dispersal estimates.

Our IBM predicts, given sufficient time, mutational differences between generalist microbes would greatly increase over space (Figure 2) following a significant decrease in mean home range and we hypothesize that this impacted EID formation (Figure 3). Since this hypothesized event occurred in the past, it is difficult to empirically test. However, databases exist with the locations of EIDs and previous work has correlated these locations with environmental and biodiversity variables (Jones et al., 2008). We therefore hypothesize that if decreased dispersal is related to zoonotic emergent disease, then adding a dispersal variable should improve the prediction of EIDs. More broadly, we hypothesize that information from the ecological past is relevant to predictions of future potential outbreaks.

The prediction of the spatial distribution of 145 zoonotic EID outbreaks was significantly improved by including the global loss of microbial dispersal following the extinctions (adding the change in fecal diffusivity (Δ*FD)* improved pseudo r^2^ by 8% (from 0.185 (null model) to 0.201 (model *FD*) while reducing AIC by ∼3% - Table 2). We started with 16 variables described in Table S1, including our six maps from Figure 1, but the model that best predicted EIDs included *ΔFD* plus reporting bias (estimated as log (Journal of Infectious Disease articles (*JID*)); richer countries with more scientists will find more EIDs), human population density and rainfall. Both change in ecotoparisite dispersal (Δ*MHR – Equation 2)* and change in fecal diffusivity *(ΔFD-Equation 7)* were highly significant but had collinearity issues (VIF>10) and we chose to include just *ΔFD* versus Δ*MHR* because it reduced AIC by a greater amount (Table 2 – model HR). Jones *et al*. 2008 found current species richness to be a significant explanatory variable because host diversity is strongly positively correlated with pathogen diversity, and we also found this on its own (Table 2). However, adding *SR*_*current*_ to our best model increases AIC (Table 2 – model SR) and we did not include it in our final model. We tested whether transmission mode (vector-borne versus non-vector-borne transmission) impacted our results and found adding either *ΔFD* and Δ*MHR* improved model performance (reduced AIC) in both models predicting vector-borne and non-vector-borne EIDs, (SI Appendix, Table S3 and S4). In a sensitivity study (SI Appendix, Table S5 and S6) we tested the resilience of our results and found that our model results remained significant under a wide range of scenarios. For instance, moving the EID location randomly by one pixel to estimate the great uncertainty in knowing the exact EID emergence coordinates, did not greatly change our results.

We then create a new EID prediction map based on model FD (Table 2 and Figure 4 top) accounting for sampling bias by removing the log(*JID*) parameter. There are many similarities of this map to the original Jones et al. (2008) map with large hotspots in regions of large human population densities. However, our map shows a more pronounced peak in southern South America and the north-eastern North America due to the impact of the Δ*FD* variable. We also estimate how the extinctions impacted EID occurrence (Figure 4 bottom) by subtracting Δ*FD* from our best EID prediction map (Figure 4 top) since if the extinctions had not happened, there would be no change in *FD* and removing this variable creates an EID prediction map with no extinctions. Without extinctions, global predicted EIDs are reduced by 24-42% (under low (0.0019) and high (0.0033) Δ*FD* scenarios since the slope of Δ*FD =* 0.0026 ± 0.0007, Table 2 – model FD). If we conservatively estimate that our model only captures about a fifth of the EID variance (total model r^2^ is highly dependent on the number of control points chosen though), then the extinctions increased EID occurrence by 5-8% (or 7-12 of the 145 total EIDs). We see the most profound differences in southern South America, eastern USA and central Eurasia where there were the most drastic decreases in body size. Using Δ*MHR* (Table 2 – model HR) in place of Δ*FD* gives similar results and global EIDs are reduced by 20-38% (under low (0.0012) and high (0.0022) Δ*MHR* scenarios since the slope *=* 0.0017+0.0005, Table 2 – model HR).

**Figure 4.**
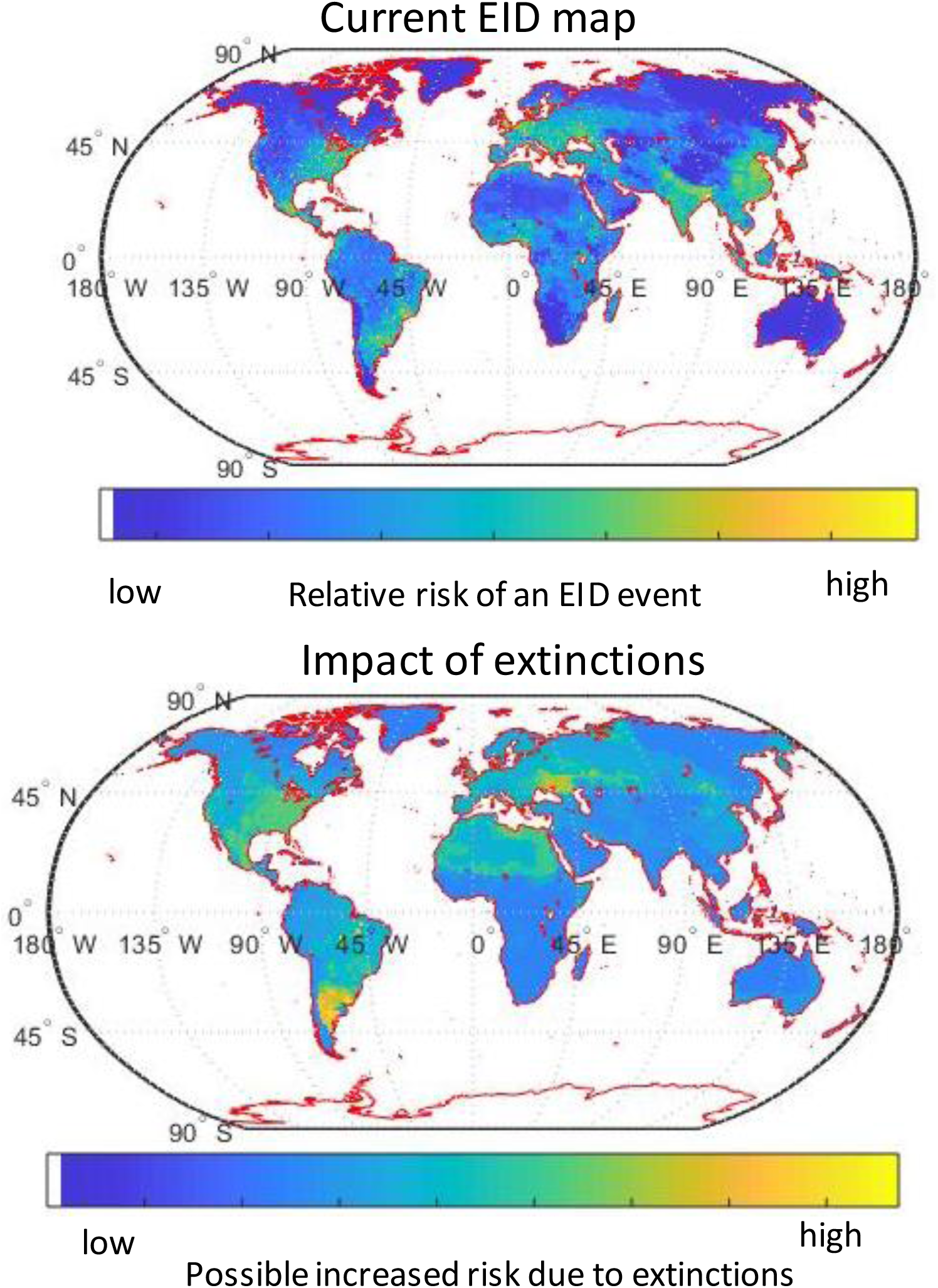
(top) A map of EID likely occurrence based on coefficients from Table 2 – model FD, but removing reporting bias (by excluding the variable log(JID)). (bottom) The current EID occurrence map (top) minus a map produced where the megafauna never went extinct (the variable Δ FD is zero). Therefore, this is a map showing where EID occurrence has increased in probability due to the megafauna extinctions. In other words, it shows “cost of extinctions” for humanity in terms of increased EID likelihood.

## Discussion

The Late Pleistocene megafauna extinctions reduced dispersal of generalist fecal microbes and ectoparasites to ∼15% of dispersal prior to the extinctions (Table 1). In certain regions such as southern South America, pathogens need >70 times as long to interact over the same distance today compared to a non-extinction world (Figure 1). A simple mechanistic model showed an order of magnitude reduction in dispersal could increase maximum mutations 4-fold (2-4 fold) (Figure 2 and SI Appendix, Table S7). We hypothesize that this pathogen speciation could have impacted EID formation through a mechanism similar to the theory of island biogeography (Reperant, 2010)(Heaney, 2000) where pathogen speciation and hence endemism increase with isolation as global dispersal distances decrease (Figure 3). In addition, this increased time in pathogen flow may impact EID formation by increasing the immune-naivety of host species because they will no longer regularly interact with as many pathogens. Our theory is supported because the change in fecal diffusivity significantly improves the prediction of EID formation (Table 2 and Figure 4). We acknowledge that changes to the range of historic mammals only explains a small amount of the variance in EID location. Many variables confound our analysis and would have been included had there been a comprehensive global dataset, such as: no past global animal abundance data, inclusion of all vector EIDs including non-generalists, not including bats, uncertainty in true EID origin.

However, despite the many problems with the EID data, the correlation of decreased past dispersal with EID formation suggests that a fraction (8%) of the variance of EID formation can be explained by an event 10-15,000 years ago. Therefore, we further explore this idea below, while acknowledging much further research is needed to empirically support it. In our IBM (Figure 2), total unique mutations saturated at ∼2,000 only after 10,000 model years, suggesting such a timeframe is reasonable for microbial evolution (supplementary methods). However, here we are suggesting that the loss of dispersal is correlated with modern disease emergence (we suggest they are also related to older diseases, such as the origin of smallpox, but we do not know the specific time or location of the emergence of these diseases into a human population, and hence, these emergences were excluded from the Jones et al (2008) dataset. Could something that happened >10,000 ybp affect diseases since 1940? Our statistical model indicates that high human population densities are necessary for EID formation (Table 2). However, high human and domestic animal densities had not arrived until the late Holocene in much of the world and closer to the timeframe of the EID database. Therefore, the early Holocene may have been a time where reduced dispersal enabled microbial speciation but these new strains of microbes did not cause EID until other elements necessary, such as high human and domestic animal population densities, were also present.

Part of the temporal discrepancy may also be due to pathogens first jumping to domesticated animals. Many domestic animals are artiodactyls similar to many of the extinct megafauna, and ungulates are the dominant hosts of zoonotic diseases (Han, Kramer, & Drake, 2016), making the evolutionary jump for pathogens from the extinct megafauna to domestic animals easier (the phylogenetic hypothesis). Domestic animals have taken over some of the functional roles of the now extinct megafauna, including their biomass (Barnosky, 2008), and it is conceivable that they also eventually hosted their pathogens, which later spread to people through close contact (Klous et al 2016, Jones et al 2011). Inbreeding of domestic animals may have reduced their natural immune defences towards parasites and pathogens (Smallbone et al., 2016). Domestic animals could also act as amplifier hosts (e.g. the Australian outbreak of Hendra virus via horses in the 90s (Mendez et al 2012)). Parasites that once attacked the now extinct megafauna, may have preferred domestic animals with limited defences to wild animals with evolved defence mechanisms. For instance, the common vampire bat *(Desmodus rotundus)* carries several blood transmitted diseases (including rabies), which feeds on domestic animals and humans, especially where populations of large animals are depleted (Bobrowiec, 2015) Wild fauna, for example, brocket deer *(Mazama* spp.) have evolved vigorous avoidance behaviour towards vampire bats (Galetti, et al., 2016). Further empirical support of this theory could come from correlating *domestic* animal EIDs to loss of past dispersal.

Are our correlations between animal extinctions and EID formation due to ecological fallacy, where correlated data are related yet not mechanistic? For example, the areas that lost the most animal species may also have had significant environmental changes that actually led to the EID formation. However, megafauna are keystone animals and ecosystem engineers and besides reducing pathogen dispersal, the loss of the megafauna ecosystem engineers would lead to drastic continental changes in ecosystem structure (Doughty et al 2016). It has been shown that recent losses of African megafauna have increased rodent-borne disease, partially because changed landscape structure following the removal of ecosystem engineers as species like elephants create better habitat for rodent and pathogen populations (Young et al 2014). Another study found total tick abundance (not species richness) increased in East Africa by 170% when herbivores >1000 kg were excluded, and by 360% with all large wildlife excluded (Titcomb *et al* 2017). Accordingly, such changes may impact EID formation.

The number of parasite species colonizing mammals scales with body size (Esser et al., 2016) and mass scaling relationships suggest that the largest extinct megafauna species would have hosted a wide diversity of tick species(Esser et al., 2016)(Galetti et al 2018). It is uncertain whether ticks would have gone extinct following the megafauna extinctions, or switched hosts (e.g., there are 63 endangered tick species associated with threatened mammals (Mihalca, Gherman, & Cozma, 2011)). This is potentially the major source of uncertainty in our hypothesis – whether past pathogens went extinct following the loss of their host or evolved to new hosts. In the future, this could potentially be tested through metagenomics to understand the range of pathogens present in deposits of extinct mammal dung.

If our theory finds further empirical support, then could we manage our ecosystems in the future to reduce infectious disease outbreaks? Would increasing dispersal capacity (either with more large animals or improved dispersal corridors) reduce EID occurrence? If increased species immune-naiveté drove increased EID occurrence, then increasing the dispersal capacity of our ecosystems would help through, for example, the conversion of fenced pasture monocultures to free-range pastoralism (Poschlod & Bonn 1998). However, if the extinctions influenced pathogen evolution, this evolution has already occurred and increasing dispersal may not help (although it could reverse the trend).

We do not discount the importance of other more recent dispersal events such as colonialism, the slave trade, or tire exportation as more recent causes of EID hotspots. Nor do we suggest that megafaunal extinctions are the sole cause of new EIDs since other causes such as human hunting behavior, human health care and GDP, land use change, density of humans/livestock (Allan et al 2017) have also been shown to impact EIDs. Our results simply suggest that a more ancient large-scale change in dispersal patterns may also have had an impact. Biodiversity has been shown to provide many ecosystem services, including disease regulation (Cunningham, Daszak, & Wood, 2017). Here, we suggest that past animal size and dispersal capacity should also be considered in understanding disease emergence. Large animals are (and always have been) most vulnerable to anthropogenic extinction pressure (Dirzo et al., 2014), and our research suggests an important step in disease regulation would be to stop current large-animal extirpations.

## Acknowledgements

We acknowledge support for this work from a NASA biodiversity grant (#)

**Source Code –** All data and code can be currently found in dropbox folder **<u>(link</u>)**

## Notes

#### Summary of Updates

This paper has now been accepted in the journal Ecography and has a digital object identifier (DOI) of: 10.1111/ecog.05209

## References

Allen, T., Murray, K. A., Zambrana-Torrelio, C., Morse, S. S., Rondinini, C., Di Marco, M., … Daszak, P. (2017). Global hotspots and correlates of emerging zoonotic diseases. Nature Communications, 8(1), 1124. doi:10.1038/s41467-017-00923-8

Andrew P. Dobson, & E. Robin Carper. (1996). Infectious diseases and human population history. BioScience, 46(2), 115–126. doi:10.2307/1312814

Barnosky, A. D. (2008). Colloquium paper: Megafauna biomass tradeoff as a driver of quaternary and future extinctions. Proceedings of the National Academy of Sciences of the United States of America, 105 Suppl 1, 11543–11548. doi:10.1073/pnas.0801918105 [doi]

Bivand, R., Hauke, J., & Kossowski, T. (2013). Computing the jacobian in gaussian spatial autoregressive models: An illustrated comparison of available methods. Geographical Analysis, 45(2), 150–179. doi:10.1111/gean.12008

Bobrowiec P. E. D. (2015). Prey preference of the common vampire bat (desmodus rotundus, chiroptera) using molecular analysis.. Journal of Mammalogy, 96(857), 54–63.

Carbone C, Cowlishaw G, Isaac NJB, Rowcliffe JM (2005) How far do animals go? Determinants of day range in mammals. Am Nat 165: 290–297.

Cunningham, A. A., Daszak, P., & Wood, J. L. N. (2017). One health, emerging infectious diseases and wildlife: Two decades of progress? Philosophical Transactions of the Royal Society of London.Series B, Biological Sciences, 372(1725), 10.1098/rstb.2016.0167. doi:20160167 [pii]

Dirzo, R., Young, H. S., Galetti, M., Ceballos, G., Isaac, N. J., & Collen, B. (2014). Defaunation in the anthropocene. Science (New York, N.Y.), 345(6195), 401–406. doi:10.1126/science.1251817 [doi]

Dormann, C. F., McPherson, J. M., Araújo, M. J., & et al. (2007). Methods to account for spatial autocorrelation in the analysis of species distributional data: A review. Ecography, 30(5) doi:10.1111/j.2007.0906-7590.05171.x

Doughty, C. E., Roman, J., Faurby, S., Wolf, A., Haque, A., Bakker, E. S., … Svenning, J. (2016). Global nutrient transport in a world of giants. Proc Natl Acad Sci USA, 113(4), 868. Retrieved from http://www.pnas.org/content/113/4/868.abstract

Doughty CE, Faurby S & Svenning JC (2016) The impact of the megafauna extinctions on savanna woody cover in south america. Ecography 38

Esser, H. J., Foley, J. E., Bongers, F., Herre, E. A., Miller, M. J., Prins, H. H., & Jansen, P. A. (2016). Host body size and the diversity of tick assemblages on neotropical vertebrates. International Journal for Parasitology.Parasites and Wildlife, 5(3), 295–304. doi:10.1016/j.ijppaw.2016.10.001 [doi]

Esser, H. J., Foley, J. E., Bongers, F., Herre, E. A., Miller, M. J., Prins, H. H. T., & Jansen, P. A. (2016). Host body size and the diversity of tick assemblages on neotropical vertebrates. International Journal for Parasitology: Parasites and Wildlife, 5(3), 295–304. doi://doi.org/10.1016/j.ijppaw.2016.10.001

Faurby, S., & Svenning, J. C. (2015a). Historic and prehistoric human-driven extinctions have reshaped global mammal diversity patterns. Divers. Distrib, 21(10) doi:10.1111/ddi.12369

Faurby, S., & Svenning, J. C. (2015b). A species-level phylogeny of all extant and late quaternary extinct mammals using a novel heuristic-hierarchical bayesian approach. Molecular Phylogenetics and Evolution, 84, 14–26. doi:10.1016/j.ympev.2014.11.001 [doi]

Faurby, S., & Svenning, J. C. (2016). Resurrection of the island rule: Human-driven extinctions have obscured a basic evolutionary pattern. The American Naturalist, 187(6), 812–820. doi:10.1086/686268 [doi]

Galetti, M., Pedrosa, F., Keuroghlian, A., & Sazima, I. (2016). Liquid lunch– vampire bats feed on invasive feral pigs and other ungulates.. Frontiers in Ecology and the Environment, 14, 505.

Galetti, M., Moleón, M., Jordano, P., Pires, M. M., Guimarães, P. R., Pape, T., … Svenning, J. (2018). Ecological and evolutionary legacy of megafauna extinctions. Biological Reviews, 93(2), 845–862. doi:10.1111/brv.12374

Han, B. A., Kramer, A. M., & Drake, J. M. (2016). Global patterns of zoonotic disease in mammals. Trends in Parasitology, 32(7), 565–577. doi:10.1016/j.pt.2016.04.007 [doi]

Heaney, L. R. (2000). Dynamic disequilibrium: A long-term, large-scale perspective on the equilibrium model of island biogeography. Global Ecology and Biogeography, 9, 59–74.

Hoppe, K. A., Koch, P. L., Carlson, R. W., & Webb, S. D. (1999). Tracking mammoths and mastodons: Reconstruction of migratory behavior using strontium isotope ratios. Geology, 27(5), 439–442.

Jones, K. E., Patel, N. G., Levy, M. A., Storeygard, A., Balk, D., Gittleman, J. L., & Daszak, P. (2008). Global trends in emerging infectious diseases. Nature, 451(7181), 990–993. doi:10.1038/nature06536 [doi]

Jones, Bethan I, D. J. Mckeever, Delia Grace, Dirk U. Pfeiffer, Florence Mutua, Jemimah Njuki, John J. Mcdermott, Jonathan Rushton, Mohammed Yahya Said, Polly J. Ericksen, Richard Kock and Silvia Ibáñez Alonso. “Zoonoses (Project 1): Wildlife/domestic livestock interactions.” (2011).

Keesing, F., Belden, L. K., Daszak, P., Dobson, A., Harvell, C. D., Holt, R. D., … Ostfeld, R. S. (2010). Impacts of biodiversity on the emergence and transmission of infectious diseases. Nature, 468(7324), 647–652. doi:10.1038/nature09575 [doi]

Kelt, D.A. and Van Vuren, D.H., (2001) The Ecology and Macroecology of Mammalian Home Range Area vol. 157, no. 6 the american naturalist

Klous, Gijs, Anke Huss, Dick J.J. Heederik, Roel A. Coutinho, (2016) Human–livestock contacts and their relationship to transmission of zoonotic pathogens, a systematic review of literature,One Health, Volume 2, 2016, Pages 65-76, ISSN 2352-7714, https://doi.org/10.1016/j.onehlt.2016.03.001.

Malhi, Y., Doughty, C. E., Galetti, M., Smith, F. A., Svenning, J. C., & Terborgh, J. W. (2016). Megafauna and ecosystem function from the pleistocene to the anthropocene. Proceedings of the National Academy of Sciences of the United States of America, 113(4), 838–846. doi:10.1073/pnas.1502540113 [doi]

McConnell, S. M., & Zavanda, M. S. (2013). The occurrence of an abdominal fauna in an articulated tapir (*tapirus polkensis*) from the late miocene gray fossil site, northeastern tennessee. Integrative Zoology, 8 doi:10.1111/j.1749-4877.2012.00320.x

Mendez DH, Judd J, Speare R. Unexpected result of Hendra virus outbreaks for veterinarians, Queensland, Australia. Emerg Infect Dis. 2012;18(1):83–85. doi:10.3201/eid1801.111006

Mihalca, A. D., Gherman, C. M., & Cozma, V. (2011). Coendangered hard-ticks: Threatened or threatening? Parasites & Vectors, 4, 71. doi:10.1186/1756-3305-4-71 [doi]

Naimi, B., Hamm, N. A. S., Groen, T. A., Skidmore, A. K., & Toxopeus, A. G. (2014). Where is positional uncertainty a problem for species distribution modelling? Ecography, 37(2), 191–203.

Poschlod P & Bonn S (1998) Changing dispersal processes in the central European landscape since the last ice age: An explanation for the actual decrease of plant species richness in different habitats?. Acta Botanica Neerlandica 47: 27–44

Reperant, L. A. (2010). Applying the theory of island biogeography to emerging pathogens: Toward predicting the sources of future emerging zoonotic and vector-borne diseases. Vector Borne and Zoonotic Diseases (Larchmont, N.Y.), 10(2), 105–110. doi:10.1089/vbz.2008.0208 [doi]

Sandom, C., Faurby, S., Sandel, B., & Svenning, J. C. (2014). Global late quaternary megafauna extinctions linked to humans, not climate change. Proceedings.Biological Sciences, 281(1787), 10.1098/rspb.2013.3254. doi: 10.1098/rspb.2013.3254 [doi]

Smallbone, W., van Oosterhout, C., & Cable, J. (2016). The effects of inbreeding on disease susceptibility: Gyrodactylus turnbulli infection of guppies, poecilia reticulata. Experimental Parasitology, 167, 32–37. doi:10.1016/j.exppara.2016.04.018 [doi]

Schmidt, K. A. & Ostfeld, R. S. Biodiversity and the dilution effect in disease ecology. Ecology (2001). doi:10.1890/0012-9658(2001)082[0609:BATDEI]2.0.CO;2

Titcomb G, et al (2017) Interacting effects of wildlife loss and climate on ticks and tick-borne disease. Proceedings of the Royal Society B 284(1862)

Tucker, M. A., Böhning-Gaese, K., Fagan, W. F., Fryxell, J. M., Van Moorter, B., Alberts, S. C., … Mueller, T. (2018). Moving in the anthropocene: Global reductions in terrestrial mammalian movements. Science, 359(6374), 466.

Wolf, A., Doughty, C. E., & Malhi, Y. (2013). Lateral diffusion of nutrients by mammalian herbivores in terrestrial ecosystems. PloS One, 8(8), e71352. doi:10.1371/journal.pone.0071352 [doi]

Wolfe, N. D., Dunavan, C. P., & Diamond, J. (2007). Origins of major human infectious diseases. Nature, 447, 279. Retrieved from http://dx.doi.org/10.1038/nature05775

Wood, C. L., McInturff, A., Young, H. S., Kim, D., & Lafferty, K. D. (2017). Human infectious disease burdens decrease with urbanization but not with biodiversity. Philos Trans R Soc Lond B Biol Sci, 372(1722)

Young HS, et al (2014) Declines in large wildlife increase landscape-level prevalence of rodentborne disease in Africa. Proc Natl Acad Sci U S A 111(19): 7036–7041.

