## Supplementary material for "Megafauna decline have reduced pathogen dispersal which may have increased emergent infectious diseases"

**Supplementary Information:**

Supplementary methods

Extended data - Figures S1-S5

Extended data - Tables S1 to S7

### Supplementary methods

### Individual based model

How might our genetic counter in our IBM translate into actual microbial speciation? We argue that it measures the accumulation of mutations in a pathogen strain that is unique to a single host. The counter does not represent mutations in general and is not able to track a specific mutation. Prior to model construction, we reasoned that reduced host dispersal would increase microbe isolation, leading to the increased divergence between microbes of different geographies. By assuming a maximal rate of microbe transfer between hosts (sharing all microbes during an interaction and resetting the counter to zero), we are minimizing the probability of individual-host-specific strains. Kishimoto et al (2015) have argued that the number of neutral mutations fixed in a population per-generation is equal to the spontaneous neutral mutation rate at the genome level (average number of neutral mutations per genome, per generation) Kishimoto, T. *et al* (2015). Like previous studies, Kishimoto et al demonstrated an *E. coli* mutation rate of approximately $1x10^{-3}$ per genome per generation Lee, et al. (2012). Therefore, we assume that $1x10^{-3}$ mutations are fixed per generation. An additional simplification of this model is that each individual animal hosts a monoclonal population of E. coli. Significant variation has been recorded in the generation times of *E. coli* (Lee, et al. (2012), Souza, et al(2002), Licht et al. (1999), Massot, *et al.* (2017), Poulsen *et al.* (1994), (Sprouffske, et al (2018)) from 30 minutes on nutrient-rich laboratory cultures Sprouffske, et al (2018), to Savageau's (1983) estimate of 40 hours in the mammalian gut (Chib et al (2017), Savageau, (1983)). Given the year-long increments of the IBM, these extremes in doubling times correspond to 17532 and 219 *E. coli* generations, respectively, or 18 and 0.2 *single nucleotide polymorphisms*. How many individual microbial mutations are within the large animal home range? In other words, how many mutation rates are equal to one model time step?

We can very roughly estimate this number with the following methodology. We estimate the home range of mammals with the following equation where M is mass (kg) (Wolf et al 2014):

***Equation S1*** - mammal herbivore home range - $0.04*M^{1.09}$ units of $km^2$

Therefore, an extinct 4000 kg mammal might have a 350 $km^2$ (±250km2) home range. We can use equation 2, originally from Damuth (1987) to estimate that within this 350 $km^2$ home range, there could live ~200,000 (±15,000), 0.1 kg mammals.

***Equation S2*** – mammal herbivore pop density - 87*M^-0.72 - units of ind/$km^2$

If each animal had an average of one tick, then there would be 200,000 ticks per large home range. If microbe populations homogenize within each tick, then we estimate 200,000 individual microbial populations with 0.2 to 18 mutations per year or 0.04e6 - 3.6e6 mutations per large home range per time step.

It is generally accepted that a 3% genomic change would constitute two distinct bacterial species. The difference in protein-coding non-housekeeping genes may be even lower. Palyas et al (1997) showed that the mean sequence divergence between closely related bacterial species and sub-species is around 1% (Palys, et al (1997), Cohan (2001)). This divergence was almost always >2 times greater than the within taxa. If the average length of a microbial genome is 3 million base pairs, then a 1-3% change would be 30,000-90,000 changes. Our model simulates that after 10,000 time steps (years), there were microbes that were able to evolve independently for >2000 time steps, which is (even in our lower bound estimate - 0.04e6 *2000>90,000) more than enough for many potential speciations (3% change) to occur.

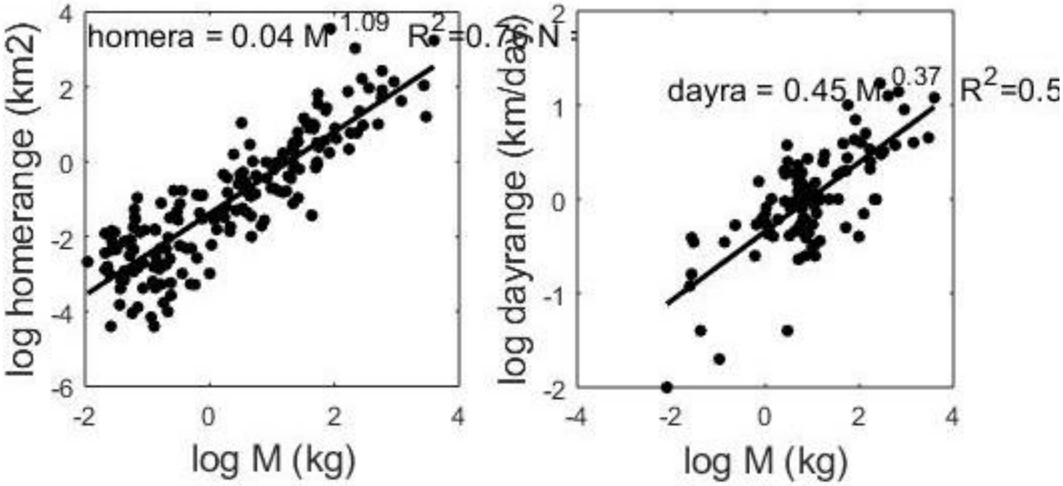


**Figure S1** – Calculation of log10 mammal herbivore mass (kg) versus log10 home range (left) and log10 day range (right)

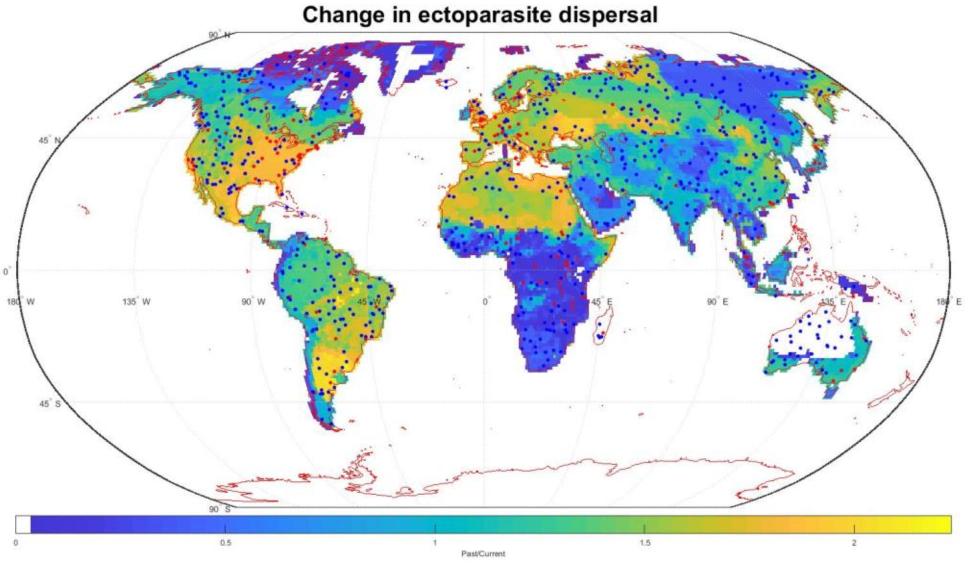


**Figure S2** – Map of zoonotic EIDs (red) and control points (blue) with log10 Δ MHR as the background.

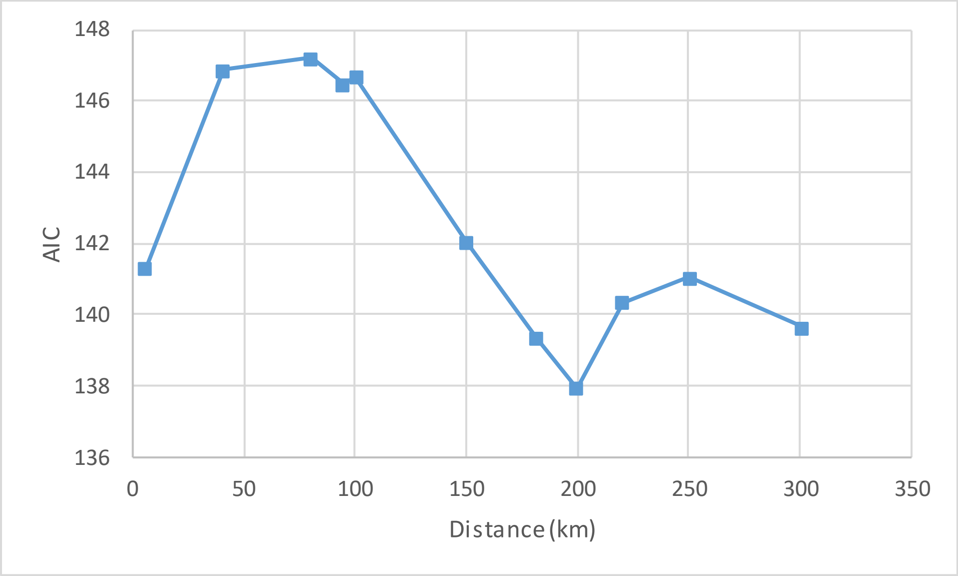


**Figure S3** – Average for 16 simulations showing a reduced AIC value with a neighborhood distance of 200km.

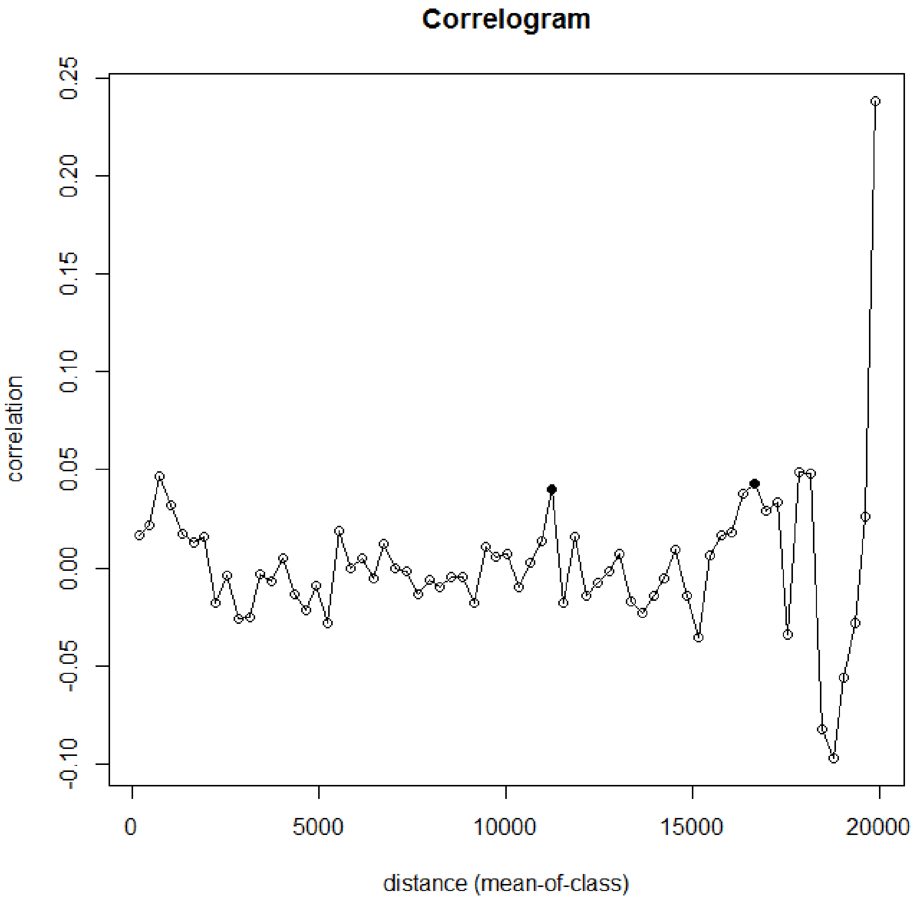


**Figure S4** – Example Correlogram for model 2. Mean of 16 correlograms (not shown) had a peak correlation of 0.058±0.02 (sd) within a distance of 5000 (mean of class).

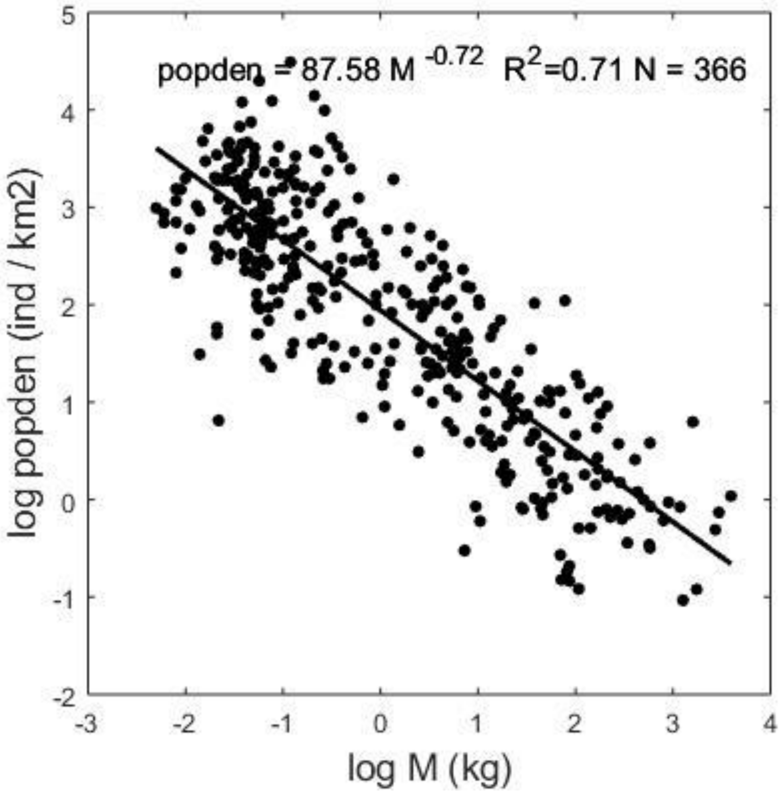


**Figure S5** – Herbivore mass versus population density.

**Table S1** – Description of our 16 variables of interest.

| Variable | Description | Reference |
|---|---|---|
| **Log(JID)** | Number of journal articles published in the Journal of Infectious Disease from Jones et al. 2008 averaged on a country basis. | Jones et al. 2008 |
| **Pop density** | Pixel level current estimates of population density based on population data from the World Bank. | The World Bank |
| **Pop change** | Country level world bank population estimates subtracting population estimates for 2015 from 2005. From http://data.worldbank.org/indicator/SP.POP.TOTL | The World Bank |
| **Δ MHR** | The change in mean home range (eq. 2) calculated by subtracting past mean home range from current mean home range. | Faurby and Svenning, 2015, Wolf et al 2014 |
| **Δ FD** | The change in mean fecal diffusivity (eq. 7) calculated by subtracting past mean fecal diffusivity from current. | Faurby and Svenning, 2015, Wolf et al 2014 |
| **$MHR_{past}$** | Past mean home range calculated as the mean home range (eq. 2) for current mammals using the IUCN species range maps and past mammals based on Faurby and Svenning (2015). | Faurby and Svenning, 2015, Wolf et al 2014 |
| **$FD_{past}$** | Past mean fecal diffusivity (eq. 7) calculated as the mean fecal diffusivity for current mammals using the IUCN species range maps and past mammals based on Faurby and Svenning (2015). | Faurby and Svenning, 2015, Wolf et al 2014 |
| **$MHR_{current}$** | Current mean home range (eq. 2) calculated as the mean home range for current mammals using the IUCN species range maps. | Faurby and Svenning, |

| | | |
|---|---|---|
| | | 2015, Wolf et al 2014 |
| **$FD_{current}$** | Current mean fecal diffusivity (eq. 7) calculated as the mean fecal diffusivity for current mammals using the IUCN species range maps. | Faurby and Svenning, 2015, Wolf et al 2014 |
| **Rain** | We used GPCC Precipitation Monitoring V4 (1.0x1.0) (180 by 360 by 120) for the period 2007 to present, based on quality-controlled data from 7,000 stations and found at http://www.esrl.noaa.gov/psd/data/gridded/data.gpcc.html. For each pixel, we found the mean monthly rainfall. | NOAA |
| **Seasonality** | For each pixel, we subtracted the difference between maximum and minimum monthly precipitation from the above reference | NOAA |
| **$SR_{current}$** | Current species richness is determined by summing up the total number of mammal species in each pixel based on the IUCN red list. | IUCN |
| **$SR_{past}$** | Past species richness sums up all mammal species per pixel based on both the IUCN red list and Faurby and Svenning (2015a). | Faurby and Svenning, 2015, |
| **Latitude** | This is the latitude of the pixel. | |
| **BWSP** | Biomass weighted species richness for current (from IUCN) and past species (from Faurby and Svenning (2016)) based on eq. 8. | Faurby and Svenning, 2015, |
| **Δ BWSP** | Past biomass weighted species richness minus current. | Faurby and Svenning, 2015, |

**Table S2** – A SAR$^{err}$ analysis to predict the presence of 145 EIDs compared with ~600 randomly generated points (generated three separate times) using the 16 following variables (further described in Table S1): *JID* -journal of infectious disease articles, human population density, change in human population density, *Δ MHR* – change in mean home range, *ΔFD* – change in fecal diffusivity, $MHR_{past}$ – MHR during the late Pleistocene, $FD_{past}$ –FD during the late Pleistocene, $MHR_{current}$ – current MHR, $FD_{current}$ current FD, Rain – average rainfall, Seasonality – difference between max and min monthly rainfall, $SR_{current}$ – Species richness during the Pleistocene, $SR_{current}$ – current species richness, latitude, *BWSP* – biomass weighted species richness and *Δ BWSP* the change (past – current) in this variable. In column one, we show the variables of interest, in column two, the mean value of each variable for all chosen points, in column three we show the individual model coefficient (value we multiply with the variable), $r^2$ and significance (*=P<*0.003125, **=* P<*06e-4, ***=* P<*06e-5*) using the Bonferroni correction to determine significance or *0.05/16= 0.003125 = *.*

| Variable | Mean value | Individual | Weighting |
|---|---|---|---|
| **(Intercept)** | | | |
| **Log(*JID*)** | 1.66 | $r^2$ = 0.046 , 0.065 *** | 0.09 |
| **Pop density** | 0.73 | $r^2$ = 0.12 , 0.14 *** | 0.08 |
| **Pop change** | 0.12 | $r^2$ = 0.000 , 0.08 | 0.01 |
| ***ΔMHR*** | 22.7 | $r^2$ = 0.035 , 0.003 *** | 0.06 |
| ***ΔFD*** | 3.05 | $r^2$ = 0.042 , 0.004 *** | 0.12 |
| $MHR_{past}$ | 9.63 | $r^2$ = 0.028 , 0.009 *** | 0.08 |
| $FD_{past}$ | 1.12 | $r^2$ = 0.038 , 0.03 *** | 0.12 |
| $MHR_{current}$ | 1.36 | $r^2$ = 0.000 , 0.0002 | 0.00 |
| $FD_{current}$ | 0.41 | $r^2$ = 0.000 , 0.002 | 0.00 |
| **Rain** | 63 | $r^2$ = 0.021 , 0.0009 ** | 0.05 |
| **Seasonality** | 109 | $r^2$ = 0.001 , 0.0001 | 0.01 |
| $SR_{current}$ | 111 | $r^2$ = 0.012 , 0.001 * | 0.13 |
| $SR_{past}$ | 123 | $r^2$ = 0.018 , 0.0012 ** | 0.16 |
| **Latitude** | 0.16 | $r^2$ = 0.002 , -0.0005 | 0.00 |
| ***BWSP*** | 13200 | $r^2$ = 0.027 , 5.89E-06 *** | 0.07 |

| | | | |
|---|---|---|---|
| ***Δ BWSP*** | 11200 | $r^2$ = 0.034 , 7.52E-06 *** | 0.07 |

**Table S3 –** Same as Table 2 in text but using only vector borne pathogens (N=51).

| Variable | Mean value | Individual | Weightings | Model 2 - pseudo r2 = 0.12±0.01 |
|---|---|---|---|---|
| (Intercept) | | | | -0.130 ± 0.060 * |
| Log(JID) | 1.66 | $r^2$= 0.020 , sl= 0.040 * | 0.067 | 0.060 ± 0.019 ** |
| Pop density | 0.60 | $r^2$= 0.075 , sl= 0.106 ** | 0.063 | 0.086 ± 0.025 ** |
| Pop change | 0.13 | $r^2$= 0.002 , sl= 0.044 | 0.006 | 0.452 ± 0.188 * |
| Δ MHR | 22.16 | $r^2$= 0.026 , sl= 0.002 * | 0.047 | 0.0019 ± 0.0010 * |
| Δ FD | 3.03 | $r^2$= 0.014 , sl= 0.024 | 0.073 | |
| $MHR_{past}$ | 10.06 | $r^2$= 0.014 , sl= 0.006 . | 0.056 | |
| $FD_{past}$ | 1.16 | $r^2$= 0.009 , sl= 0.065 | 0.075 | |
| $MHR_{current}$ | 1.72 | $r^2$= 0.000 , sl= -0.0005 | -0.001 | |
| $FD_{current}$ | 0.44 | $r^2$= 0.000 , sl= -0.0039 | -0.002 | |
| Rain | 60 | $r^2$= 0.012 , sl= 0.0007 . | 0.044 | 0.0005 ± 0.0004 |
| Seasonality | 106 | $r^2$= 0.002 , sl= 0.0002 | 0.017 | |
| $SR_{current}$ | 111 | $r^2$= 0.019 , sl= 0.0014 * | 0.150 | |
| $SR_{past}$ | 123 | $r^2$= 0.022 , sl= 0.0013 * | 0.165 | |
| Latitude | 29 | $r^2$= 0.003 , sl= -0.0006 | -0.018 | |
| BWSP | 14021 | $r^2$= 0.009 , sl= 2.59E-06 | 0.036 | |
| Δ BWSP | 11391 | $r^2$= 0.018 , sl= 5.71E-06 * | 0.065 | |

**Table S4 –** Same as Table 2 in text but using only non-vector borne pathogens (N=94).

| **Variable** | **Mean value** | **Individual** | **Weightings** | **Model 2 - pseudo r2 = 0.12±0.01** |
|---|---|---|---|---|
| (Intercept) | | | | -0.084 ± 0.023 *** |
| Log(JID) | 1.64 | $r^2$= 0.036 , sl= 0.049 *** | 0.081 | 0.052 ± 0.009 *** |
| Pop density | 0.70 | $r^2$= 0.118 , sl= 0.111 *** | 0.078 | 0.111 ± 0.013 *** |
| Pop change | 0.11 | $r^2$= 0.000 , sl= -0.010 | -0.001 | |
| Δ MHR | 25.01 | $r^2$= 0.018 , sl= 0.002 *** | 0.042 | |
| Δ FD | 3.20 | $r^2$= 0.020 , sl= 0.030 *** | 0.096 | |
| $MHR_{past}$ | 10.39 | $r^2$= 0.014 , sl= 0.005 ** | 0.051 | |
| $FD_{past}$ | 1.18 | $r^2$= 0.014 , sl= 0.069 ** | 0.081 | |
| $MHR_{current}$ | 1.50 | $r^2$= 0.000 , sl= -0.002 | -0.002 | |
| $FD_{current}$ | 0.42 | $r^2$= 0.000 , sl= -0.027 | -0.011 | |
| Rain | 63 | $r^2$= 0.038 , sl= 0.00099 *** | 0.062 | 0.00047± 0.00020 * |
| Seasonality | 111 | $r^2$= 0.001 , sl= 0.00009 | 0.010 | |
| $SR_{current}$ | 111 | $r^2$= 0.011 , sl= 0.00079 * | 0.087 | |
| $SR_{past}$ | 124 | $r^2$= 0.015 , sl= 0.00088 ** | 0.109 | |
| Latitude | 30 | $r^2$= 0.001 , sl= -0.00030 | -0.009 | |
| BWSP | 14526 | $r^2$= 0.010 , sl= 2.53E-06 * | 0.037 | |
| Δ BWSP | 12326 | $r^2$= 0.016 , sl= 4.02E-06 *** | 0.050 | |

**Table S5** – Description of the variables modified in the sensitivity analysis and a description of what the uncertainty is.

| Variable | Description | Uncertainty | Reference |
|---|---|---|---|
| **EID coordinates** | These are the coordinates of the EIDs taken from Jones et al. 2008. | These coordinates are highly uncertain as it is biased towards the nearest medical facility. To access this uncertainty, we randomly move the coordinates by one pixel to account for the fact that many such EIDs likely arose nearby but not in the nearest medical facility. | Jones et al 2008 |
| **Animal body mass** | The estimated mean body mass of a species | The uncertainty of individual species are generally captured with +/-25. However, we use an error of +/-10% which is for entire communities because as long as the estimates are unbiased the combined error for all co-occurring species will be smaller since some are under and some are over estimated. | Faurby and Svenning, 2015, Wolf et al 2014 |
| **FD coefficient** | This is the fecal diffusivity or the average distance moved by a gut pathogen in either the Late Pleistocene or today. | This mass scaling relationship is based on the day range of 113 mammal species and estimated passage time. The percent error on the slope is 9%. This error will quantify the variability in day range between species but will not quantify other potential sources of error such as the equal likelihood of any animal picking up and transporting a gut pathogen. | Faurby and Svenning, 2015, Wolf et al 2014 |
| **MHR coefficient** | This is the mean home range or the average distance moved by an ectoparasite in either the Late Pleistocene or today. | This mass scaling relationship is based on 171 mammal species. The percent error on the slope is 4%. This error will quantify the variability in home range between species but will not quantify other potential sources of | Faurby and Svenning, |

| | | | |
|---|---|---|---|
| | | error such as the equal likelihood of any animal picking up and transporting an ectoparasite. | 2015, Wolf et al 2014 |
| **Combined** | Here we combine the uncertainties (max and min) of the above three variables (body mass, FD, and MHR). | We combine the max and min values of each of the three above variables and rerun our statistical analysis under a high and low scenario. | Faurby and Svenning, 2015, Wolf et al 2014 |

**Table S6 –** Values used in our sensitivity analysis: the estimated range in uncertainty, how this uncertainty was assessed, variable tested, and the low and high estimate for that variable. Expert opinion was estimated by a group of experts (the authors of the paper) of the variable value in which the group was 95% certain the true value would fall within. If the number is calculated as a slope, then the standard deviation on the slope is the potential error.

| **Variable** | **Value used** | **Potential error estimate** | **How the error was assessed** | **Variable tested** | **Low value** | **High value** |
|---|---|---|---|---|---|---|
| **EID coordinates** | pixels | Move pixel | Expert opinion | Δ MHR | $R^2$=0.04<br>Sl = 0.003*** | |
| **Animal body mass** | M | ±30% for megafauna<br>±10% for current animals | Expert opinion | Δ BWSR | $R^2$=0.02<br>Sl = 7e-6** | $R^2$=0.025<br>Sl = 4e-6*** |
| **FD coefficient** | Eq 7 | ±9% | Slope error | Δ FD | $R^2$=0.02<br>Sl = 0.05*** | $R^2$=0.03<br>Sl = 0.03*** |
| **HR coefficient** | Eq 2 | ±4% | Slope error | Δ HR | $R^2$=0.03<br>Sl = 0.003*** | $R^2$=0.04<br>Sl = 0.002*** |
| **Combined** | Change HR, FD, and biomass | FD =±9%, HR = ±4%, ±30% for megafauna<br>±10% for current animals | See above | Δ HR | $R^2$=0.03<br>Sl = 0.003*** | $R^2$=0.03<br>Sl = 0.002*** |

**Table S7** – Sensitivity study from our IBM for the microbe at each time step with the maximum number of genetic changes for small home range divided by large home range simulations ± sd (N=3 runs). We ran all simulations with the middle value for the variable not of interest, except for time steps, which was 1000 for all simulations except time step. For example, for our 20,000-year simulation, we ran with 10% of cells occupied by both species and a grid size of 500 by 500.

| **variable** | **Value** | **High** | **Medium** | **Low** |
|---|---|---|---|---|
| Time steps | 20000/ 5000/ 1000 | **3.6 ± 0.81** | 2.3 ±**0.66** | 2.2 ± **0.85** |
| Original stock of species 1 (% of grid cells) | 5/ 10/ 20 | **3.8 ± 1.0** | **5.1 ± 1.4** | **1.8 ± 0.03** |
| Original stock of species 2 (% of grid cells) | 20/ 10/ 5 | **3.4 ± 1.9** | **2.1 ± 1.0** | **1.9 ± 1.0** |
| % of animals with pathogen | 5/ 10/ 20 | **3.3 ±0.3** | **4.5 ±1.0** | **1.9 ±0.5** |
| Size of grid | 300/500/1000 | **5.7 ± 4** | **2.7 ± 0.75** | **2.2 ± 0.48** |
| **Average** | | **4.2 ± 1** | **3.3 ± 1.6** | **1.9 ± 0.19** |

**Supplemental References**


Chib, S., Ali, F. & Seshasayee, A. S. N. Genomewide Mutational Diversity in *Escherichia coli* Population Evolving in Prolonged Stationary Phase. *mSphere* **2,** e00059-17 (2017).

Cohan, F. M. Bacterial species and speciation. *Syst. Biol.* **50,** 513–524 (2001).

Damuth J (1987) Interspecific Allometry of Population-Density in Mammals and Other Animals - the Independence of Body-Mass and Population Energy-Use. Biol J Linn Soc 31: 193–246.

Faurby S, Svenning JC. A species-level phylogeny of all extant and late Quaternary extinct mammals using a novel heuristic-hierarchical Bayesian approach. Mol Phylogenet Evol 2015 Mar;84:14-26.

Jones KE, Patel NG, Levy MA, Storeygard A, Balk D, Gittleman JL, et al. Global trends in emerging infectious diseases. Nature 2008 Feb 21;451(7181):990-993.

Kishimoto, T. *et al.* Molecular Clock of Neutral Mutations in a Fitness-Increasing Evolutionary Process. *PLoS Genet.* **11,** (2015).

Lee, H., Popodi, E., Tang, H. & Foster, P. L. Rate and molecular spectrum of spontaneous mutations in the bacterium Escherichia coli as determined by whole-genome sequencing. *Proc. Natl. Acad. Sci.* **109,** E2774–E2783 (2012).

Licht, T. R., Tolker-Nielsen, T., Holmstrøm, K., Krogfelt, K. A. & Molin, S. Inhibition of Escherichia coli precursor-16S rRNA processing by mouse intestinal contents. *Environ. Microbiol.* **1,** 23–32 (1999).

Massot, M. *et al.* Day-to-day dynamics of commensal Escherichia coli in Zimbabwean cows evidence temporal fluctuations within a host-specific population structure. *Appl. Environ. Microbiol.* **83,** (2017).

Palys, T., Nakamura, L. K. & Cohan, F. M. Discovery and Classification of Ecological Diversity in the Bacterial World: The Role of DNA Sequence Data. *Int. J. Syst. Bacteriol.* **47,** 1145–1156 (1997).

Poulsen, L. K. *et al.* Spatial distribution of Escherichia coli in the mouse large intestine inferred from rRNA in situ hybridization. *Infection and Immunity* **62,** 5191–5194 (1994).

Savageau, M. A. Escherichia coli Habitats, Cell Types, and Molecular Mechanisms of Gene Control. *Am. Nat.* **122,** 732–744 (1983).

Souza, V., Castillo, A. & Eguiarte, L. E. The Evolutionary Ecology of Escherichia coli. *Am. Sci.* **90,** 332–341 (2002).

Sprouffske, K., Aguílar-Rodríguez, J., Sniegowski, P. & Wagner, A. High mutation rates limit evolutionary adaptation in Escherichia coli. *PLoS Genet.* **14,** (2018).

Wolf A, Doughty CE, Malhi Y. Lateral diffusion of nutrients by mammalian herbivores in terrestrial ecosystems. PLoS One 2013 Aug 9;8(8):e71352.
